# Intersection of motor volumes predicts the outcome of ambush predation of larval zebrafish

**DOI:** 10.1101/626549

**Authors:** Kiran Bhattacharyya, David L. McLean, Malcolm A. MacIver

## Abstract

The escape maneuvers of animals are key determinants of their survival. Consequently these maneuvers are under intense selection pressure. Current work indicates that a number of escape maneuver parameters contribute to survival including response latency, escape speed, and direction. This work has found that the relative importance of these parameters is context dependent, suggesting that interactions between escape maneuver parameters and the predatory context together determine the likelihood of escape success. However, it is unclear how escape maneuver parameters interact to contribute to escape success across different predatory contexts. To clarify these issues, we investigated the determinants of successful escape maneuvers by analyzing the responses of larval zebrafish to the attacks of dragonfly nymphs. We found that the strongest predictor of the outcome was the time needed for the nymph to reach the fish’s initial position at the onset of the attack, measured from the time that the fish initiates its escape response. We show how this result is related to the intersection of the swept volume of the nymph’s grasping organs with the volume containing all possible escape trajectories of the fish. By analyzing the intersection of these volumes, we compute the survival benefit of recruiting the Mauthner cell, a neuron in anamniotes devoted to producing escapes. We discuss how escape maneuver parameters interact in determining escape response. The intersection of motor volume approach provides a framework that unifies the influence of many escape maneuver parameters on the likelihood of survival.

## 1 Introduction

An escalating arms race in predator-prey interactions is considered a major driving force behind the evolution of diverse animal morphologies [1] and consequently of the nervous systems controlling them [2]. Within these interactions, the escape maneuvers of prey are under significant selection pressure since they directly affect survival. The evolutionary pressure shaping escape maneuvers selects for those parameters of the maneuver and the underlying neural circuitry that contribute to evasion success [2]. Studies have suggested that response latency, speed, and direction of an escape maneuver are all relevant contributors to success [3, 4, 5].

However, studies have produced differing results about the influence of these parameters on the outcome. The optimality of a single escape direction for increasing distance [6, 7, 8] and the unpredictability of variable escape directions [9, 10, 11] are both hypothesized to increase escape success. Moreover, in many animals, escapes can be initiated with or without the activation of large diameter command neurons devoted to generating the shortest latency escapes with the fastest speeds [12, 13, 14]. Specifically, larval zebrafish (*Danio rerio*), a popular model system for linking neural circuits to escape behaviors [15], are dramatically less likely to survive an attack from a predator after photo-ablation of the command-like neurons devoted to generating escape responses in fish called Mauthner cells [16]. However, other results have both disputed and supported the utility of fast speeds for increasing evasion success [17, 18, 19, 20, 21, 22, 23, 24, 25]. Previous studies have argued that these differing results are due to the context-dependent importance of escape maneuver parameters [26, 27, 11]. This argument suggests that, beyond the influence of any single parameter, the interaction between escape maneuver parameters and the predation context is critical in determining the likelihood of escape success. However, it is unclear how escape maneuver parameters interact with each other in various predatory contexts to contribute to escape success.

To better understand the determinants of successful escapes, we have studied the predation of larval zebrafish by an ambush predator, the dragonfly nymph (*Sympetrum vicinum*). Dragonfly nymphs hunt by remaining motionless and waiting for prey to come within ambush distance [28, 29] (Fig. 1). Once in striking range, the nymphs attack prey with their hydraulically-powered prehensile labial masks [30, 31, 32, 33] which extend outward to grasp the prey with palps and confine them in a spoon-shaped bowl [34, 35], hereafter referred to as the mask. While the species of nymph considered here is native to North America, whereas zebrafish are native to South Asia. However, there are many species of dragonfly native to South Asia where they are believed to be natural predators of larval zebrafish [36]. Zebrafish are therefore very likely to have evolved to evade the strikes of the extending labial masks of the nymphs of these species, which are likely to resemble the strikes of the nymph considered here. Similarly, *Sympetrum vicinum* is a voracious predator that has evolved to feed on larval fish native to North America among other prey during its aquatic stage of life.

**Figure 1:**
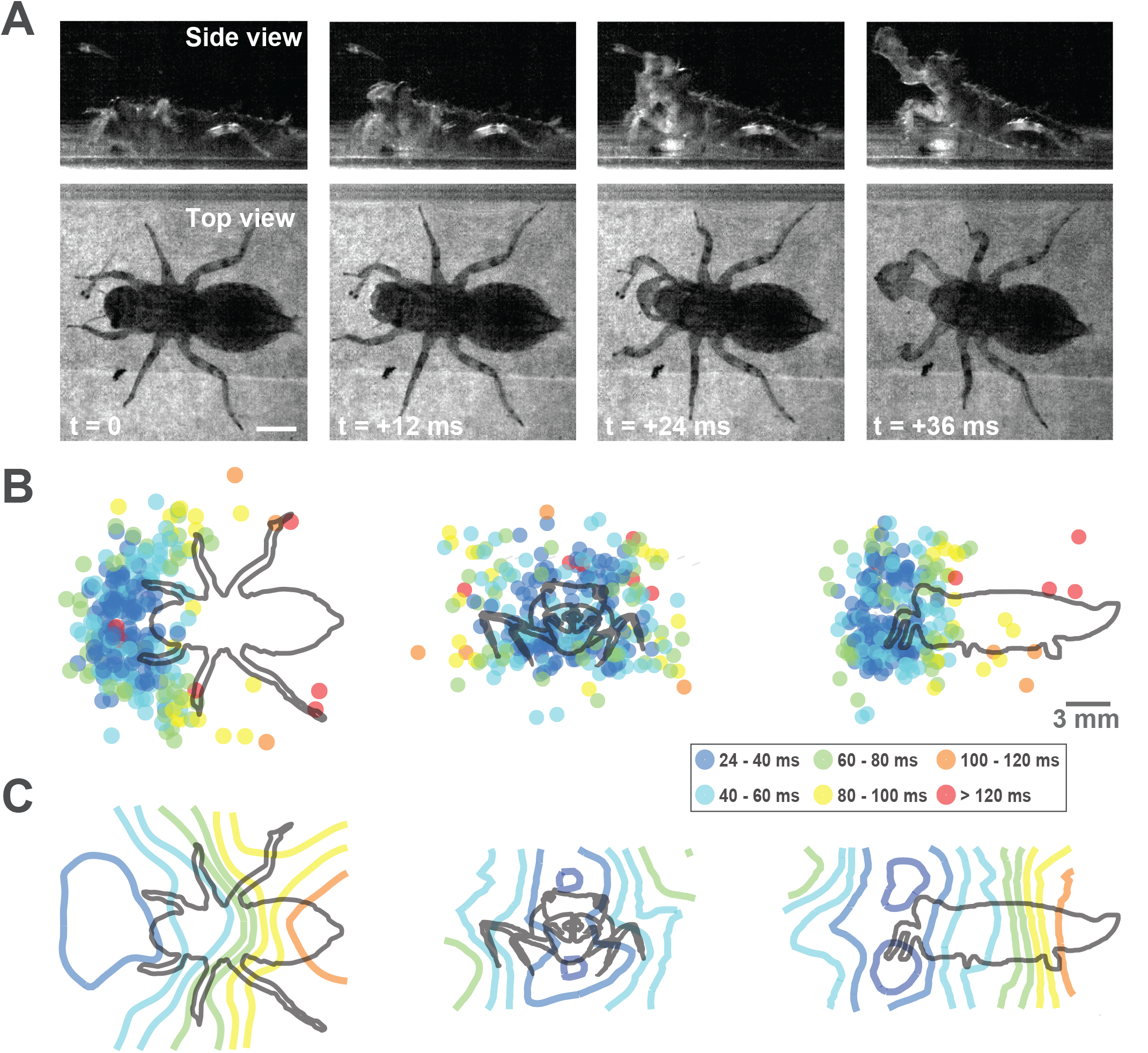
Representative dragonfly nymph strike, and prehensile mask motor volume. A) Top and side view of a strike. Scale bar: 2.5 mm. B) Top, front, and side view orthographic projections of prehensile mask strike positions colored to represent the time duration of the strike. (Number of nymphs = 5, nymph body length mean *±* std = 14.5 *±* 1.3 mm, number of attacks = 159). C) Top, front, and side view cross-sections of surfaces representing isochrones for the duration of the strike from the time of initial movement detected through high-speed videography. The 3-dimensional isochronic surfaces define the prehensile mask motor volume.

Studying responses to an ambush predator simplifies the analysis of the impact of escape maneuver parameters on survival than a similar analysis with a pursuit predator, such as adult dragonfly predatory attacks [37, 38]. This is because responses to pursuit predation involves a series of movements where it is difficult to determine which parameter ultimately led to successful escape. In contrast, ambush predation, which involves sudden strikes by predators on unsuspecting prey [39], does not provide time for a series of movements. Consequently, we focused on single escape responses of larval fish to the ballistic strikes of dragonfly nymphs to find the parameters most predictive of the outcome.

Using the time needed to extend the mask to a position in space, along with our detailed analysis of the kinematics of predator-prey encounters, we created a model of the mask motor volume—the volume swept by the grasping appendage over a given amount of time [40]. The fluid perturbations caused by the extending mask modulated the timing and kinematics of the fish escape response. Upon analysis of various parameters defining the escape response, we found that the time remaining for the mask to reach the (typically stationary) initial fish position, measured from the moment of the start of the fish’s escape response, was most predictive of escape success. Thus, if a larval fish is too close to a nymph, or if the fish is further away but too delayed in its response to the nymph attack, the time remaining for the mask to reach the fish’s position from the start of its escape response will be low, and the fish’s chance of survival will decline. Ultimately, our findings are a consequence of the intersection of the swept volume of the mask and the volume containing all possible trajectories of the fish within the time remaining to mask arrival—the fish’s motor volume [40].

We show how the interaction between the prey motor volume and the predator swept volume accounts for the influence of various escape maneuver parameters and provides a unique approach to analyze the utility of specific escape movements Additionally, we use the analysis of motor volumes to computationally estimate the survival benefit of recruiting the Mauthner cell, the giant neuron devoted to producing escape maneuvers in fish [41, 42, 43, 44]. We discuss how our conclusions generalize to other predator-prey interactions and extend the existing understanding of the selection pressure on escape responses along with their neural execution.

## 2 Materials and Methods

### 2.1 Larval zebrafish

Experiments were performed using 5–7 days-post-fertilization zebrafish larvae *Danio rerio* with body lengths of 4–5 mm, obtained from a laboratory stock of wild-type adults. At these early stages of development, zebrafish have not yet sexually differentiated. Fish in our custom-fabricated breeding facility (Aquatic Habitats, Beverly, MA, USA) were maintained at 28.5°C in system water (pH 7.3, conductivity 550 *μ*S) on a 14 hr:10 hr light:dark cycle. All recordings of behavior were performed at room temperature (24°C) water from the aquatic facility, hereafter referred to as system water. Animals were treated in accordance with the National Institutes of Health Guide for the Care and Use of Laboratory Animals and experiments were approved by the Northwestern University Institutional Animal Care and Use Committee.

### 2.2 Dragonfly nymphs

Dragonfly nymphs (*Sympetrum vicinum*) were acquired from a commercial vendor (Carolina Biological Supply Company, Burlington, NC, USA). The age of the nymphs were unknown but at this stage in development before metamorphosis, they have not yet sexually differentiated. Each nymph was stored at room temperature (24°C) in a separate tank with system water. Water in the tank was replaced weekly.

### 2.3 Behavior recordings

All recordings of behavior were performed at room temperature (24°C) system water. Five dragonfly nymphs of approximately the same size were selected (nymph body length mean *±* std = 14.5 mm *±* 1.3 mm) for all experiments since dramatic differences in size could change the size of the mask and the locomotor performance of the strike. For each experiment, a single dragonfly nymph was selected and placed into an arena within a square plastic dish (25 mm width, 100 mm length, 15 mm height, Thomas Scientific, Swedesboro, NJ, USA) with room temperature system water and allowed to acclimate for 15 minutes. The arena constrained the dragonfly nymph to move within the field of view of the dissection microscope (Stemi-2000; Carl Zeiss Microscopy, Thornwood, NY, USA). After acclimation, 1–5 larval zebrafish were introduced into the arena. More than one larval zebrafish was added to the arena in order to increase the chances of a strike from the nymph since it did not actively hunt the larva but chose to sit and wait until a larva came into striking distance. The dragonfly nymph would strike at a single zebrafish larva.

If the strike of the nymph was accurate and the larva initiated an escape response, the parameters of the fish escape responses were recorded. The strike of the dragonfly nymph was considered to be accurate if the labial mask extended to a final position *±*2 mm from where the larval fish’s swim bladder was immediately before the start of the strike. On the other hand, an inaccurate strike was one where the dragonfly nymph aimed its labial mask at a position where the fish was not present, even before the strike began. After a dragonfly nymph strike, the larvae were removed from the arena and new larvae were introduced in order to ensure that larvae were not acclimated to dragonfly nymph strikes.

To observe the dragonfly nymphs strikes and fish escape responses in our assay, videos were recorded using a high-speed camera (FASTCAM 1024 PCI; Photron, San Diego, CA, USA) attached to the microscope providing a top view into the arena. Additionally, a 100 mm long equilateral acrylic prism (Carolina Biological Supply Company, Burlington, NC, USA) was placed at the edge of the square petri dish to provide a side view perspective into the dish within the same image. The top and side views used for imaging were calibrated with black lines 15 mm in length that were viewed from both top and side views. This allowed for a pixel to mm conversion. Both views had nearly the exact same 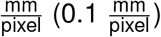 conversion suggesting that no extra corrections were necessary for the longer path length of the light for the side view perspective. The imaging set-up was not disturbed over the course of the experiments. Images were collected at 250 fps at 1X magnification.

The animals were illuminated from below, with diffused white light from the microscope for the top view. Additional illumination was provided by a halogen lamp with an articulating arm from the side for the side view. The exact amount of illumination was not measured. However, we do not believe the visual systems of the dragonfly nymph or the zebrafish larvae were compromised since they both performed naturalistic behaviors.

### 2.4 Behavior analysis

A MATLAB (MathWorks, Natick MA, USA) graphical user interface (GUI) was developed to manually track the orientation of the nymph and the position of the mask in top and side view. Tracking of the nymph was accomplished by clicking on the center of the head between the eyes followed by clicking on the position of the anus immediately before the beginning of the strike. During the strike, the center of the nymph mask was also tracked over the course of its extension. The nymph’s body and mask were tracked in top and side views. The orientation and total head yaw for the fish during the initial bend of the escape maneuver was also hand-tracked with a separate Matlab GUI in both top and side view. Larval fish were tracked by clicking on the head, the swim bladder, and the end of the tail from the start of the escape response and through the initial bend. During the propulsive stage of the escape response, the fish was tracked with an MATLAB-based automated tracker to estimate swimming velocity for the subset of escape responses that remained within the field of view. The automated tracker used thresholding and blob-detection to track the fish over the course of the escape trajectory in the top and side views. Pixel positions in the two views were then transformed into their corresponding 3-dimensional (3D) coordinates. Since the top and side view were in the same image and shared a spatial axis, combining corresponding points in the top and side views into a single 3D point involved combining points along the shared axis. Our results are presented in nymph- or fish-centered coordinate systems depending which coordinate system was more appropriate for analysis.

### 2.5 Dragonfly nymph prehensile mask motor volume model

The 3D position of each strike relative to the orientation of the dragonfly nymph body was computed with vector mathematics. Each point in the 3D point cloud of mask strike positions corresponded to a mask extension time— the time taken for the mask to reach that position in space (Figure 1). A k-nearest neighbor model [45] was trained to predict the mask extension time given a 3D position.

A k-nearest neighbor model estimates the value of a parameter at some point (here, the mask extension time for a position in 3D space) using the known value of that parameter of the k nearest neighbors to the point of interest. We tested integer values of k ranging from 1 to 10. We performed 10-fold cross validation for these integer values of k. This involved splitting the dataset into 10 parts and using 9 of the parts to predict the time needed to reach the 3D positions in the last part. This cross validation was done for each part (therefore, 10-fold) for each value of k to see which value of k provided the predictions with the lowest mean-squared error. We found that k = 5 provided the lowest overall error. This model trained with 5 nearest neighbors was used to generate the prehensile mask motor volume seen in Figure 1C.

### 2.6 Neomycin treatment of larval zebrafish

We tested the role of flow sensing in fish escape responses by compromising the lateral line in a group of larvae by exposure to a 250 *μ*mol solution of neomycin sulphate (Sigma Aldrich, St.Louis, MO) for a 30 min period, followed by a 3h recovery prior to experiments. This technique was developed in previous studies [46, 47], where it was shown to induce cell death in lateral line hair cells while leaving inner ear hair cells intact. However, we allowed for a longer recovery period after neomycin treatment to ensure recovery. After recovery, larval fish were monitored to confirm that they performed spontaneous swimming behaviors and responded with escape maneuvers to touch stimuli delivered with a tungsten filament. These larvae were then introduced into the dish with the dragonfly nymph and tested as described above.

### 2.7 Approximating fluid velocity at the fish due to mask extension

To gain further insight into the role of the perturbed fluid due to mask extension in generating the fish escape response, we used a potential flow approximation to estimate the fluid velocity at the fish due to mask extension. Mask velocity alone was not a good proxy for the perturbed fluid flow at the fish position since it does not take into account the distance of the fish from the mask. We started with an established analytical solution to the flow velocity field expressed in spherical coordinates around a sphere of radius *a* moving through an incompressible, inviscid fluid at a time dependent velocity *V* (*t*) [48].

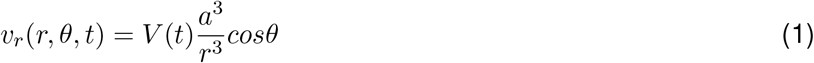

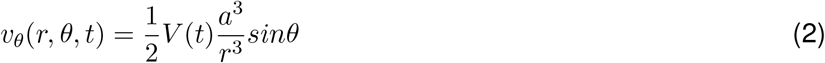

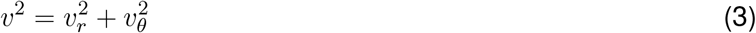

Where *v* is the flow velocity expressed in terms of the radial and angular components—*v*_*r*_ and *v*_*θ*_—for a position at some distance *r* ≥ *a* and angle *θ* from the sphere with respect to its velocity vector at some time *t*. We expect the flow pattern around the sphere to be axisymmetric around the velocity vector (i.e. independent of *φ*). In this set of equations, we can verify that when *r* → ∞ then *v* → 0, as we would expect. We used these equations with the velocity of the tracked labial mask as the value for *V* (*t*) to estimate the flow velocity *v* at the fish which was some distance *r* ≥ *a* and angle *θ*. The value of *a* was set to 1.6 mm which approximated the radius of the smallest circle enclosing the frontal projection of the mask (average diameter of 3.2 mm across all specimens).

In certain cases, the dragonfly nymph would also have to move its head during the mask extension to adjust pitch and yaw angles of the attack. This potential flow approximation does not account for this head movement. However, the head movement of the nymph is expected to contribute minimally to the already intense hydrodynamic stimulus from the extending mask, which moves a lot faster and gets progressively much closer to the fish in comparison to the head.

We used this potential flow approximation to estimate the flow velocity at the fish over other methods, such as computational fluid dynamics, due to its simplicity. Our intention was not to compute the exact magnitudes of flow velocities but rather to have relative estimates for several scenarios so that we could investigate how higher and lower flow velocities may have influenced the fish escape response differently.

### 2.8 Predicting escape outcome with each parameter

We parameterized the dragonfly nymph strike and the fish escape response to determine which parameter had the most influence on the fish escape outcome. The specific statistical tests used to determine the influence of parameters are described in the Results. To investigate the interaction of escape maneuver parameters, we trained random forest classifiers [**?**]. Random forest classifiers build an ensemble of decision trees to predict a target variable (escape success or failure) based on the values of features (attack azimuth, attack elevation, mask extension time, etc.). They do this by bootstrapping the dataset (sub-selecting from the features and the data points) to build a decision tree for each bootstrapped sample. Each decision tree generates a prediction for the target variable and the votes from the ensemble of decision trees are used to generate the final prediction of the random forest classifier. Classifiers were built with each of the following parameters to predict the binary outcome of escape failure (0) or escape success (1).

- **Attack azimuth:** the azimuth (−180° – 180°) of the dragonfly nymph strike relative to the orientation of the larval zebrafish (Figure 5B).
- **Attack elevation:** the elevation (−180° – 180°) of the dragonfly nymph strike relative to the orientation of the fish (Figure 5C).
- **Mask extension time:** the time needed to extend the mask to where the fish was located just before the initiation of an escape response (Figure 5D).
- **Bend duration:** the duration of the initial bend (Figure 4I – L) of the larval zebrafish escape response.
- **Bend velocity:** the velocity 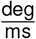 of the initial bend of the larval zebrafish escape response.
- **Response latency:** the latency of the fish escape response from the start of the nymph’s strike.
- **Time remaining at escape:** the time left from the initiation of the fish escape response until the mask reaches the initial position of the fish

**Figure 2:**
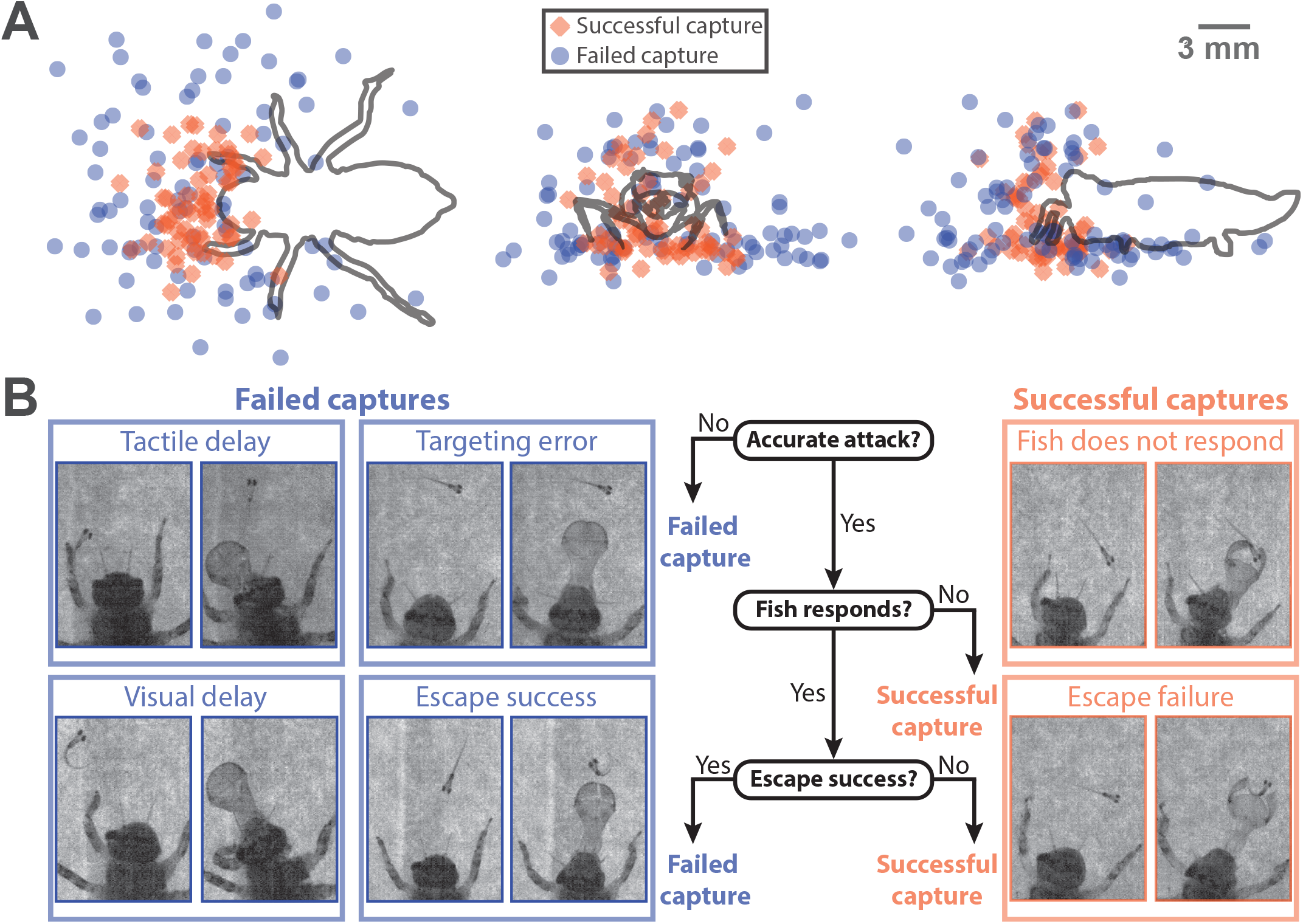
Causes of successful or failed captures by the nymph. A) Top (x-y), front (y-z), and side (x-z) view orthographic projections of 3D initial fish positions before the start of the nymph’s strike with points colored to represent the outcome of a successful or failed capture by the nymph. B) Process diagram demonstrating the sequence of events and causes of a successful or failed capture by the nymph with representative examples of each branching event in the process.

**Figure 3:**
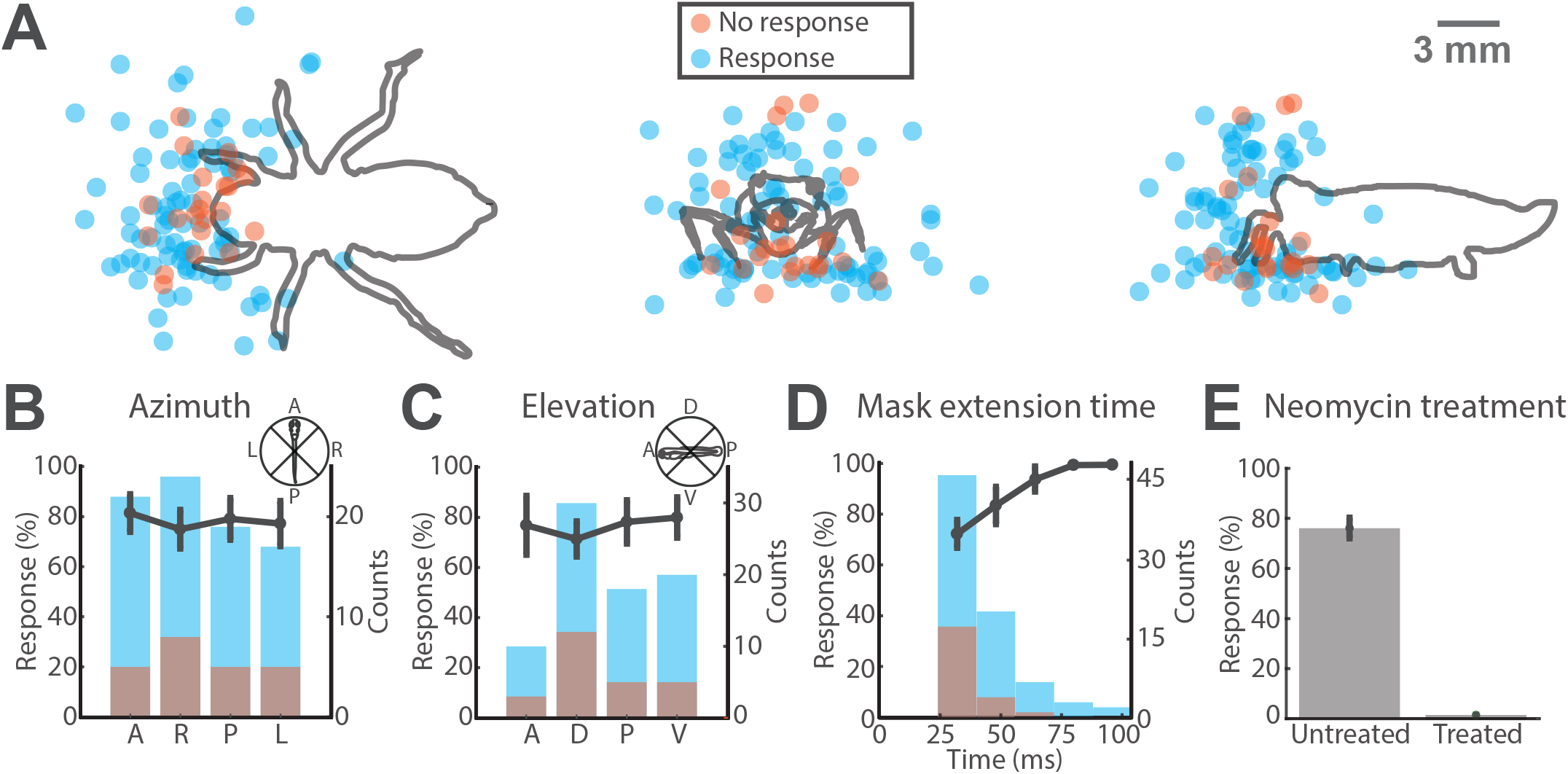
Likelihood of larval zebrafish response to an accurate strike. A) Top (x-y), front (y-z), and side (x-z) view orthographic projections of 3D initial fish positions before the start of an accurate strike with points colored to represent whether the fish responded. B–D) Heights of overlapping, partially transparent bars represent the count of strikes within that bin (right y-axis) and the color represents whether the fish responded or not. The line plot shows the mean *±* sem fish response probability for each bin (left y-axis, number of nymphs = 5 for all points with error bars). B) Fish response probability based on the azimuthal position of the nymph head with respect to the fish. There are no significant differences between response probabilities for each azimuthal quadrant. C) Fish response probability based on the elevation of the nymph head with respect to the fish. There are no significant differences between response probabilities for each elevation quadrant. D) Fish response probability based on the mask extension time to the fish position. The fish is significantly more likely to respond for longer extension times. E) Compares overall response probability of neomycin treated fish with untreated fish (untreated fish: number of trials = 109, untreated fish: number of nymphs = 5, treated fish: number of trails = 61, treated fish: number of nymphs = 4). Neomycin treated fish are far less likely to respond to a strike than untreated fish.

**Figure 4:**
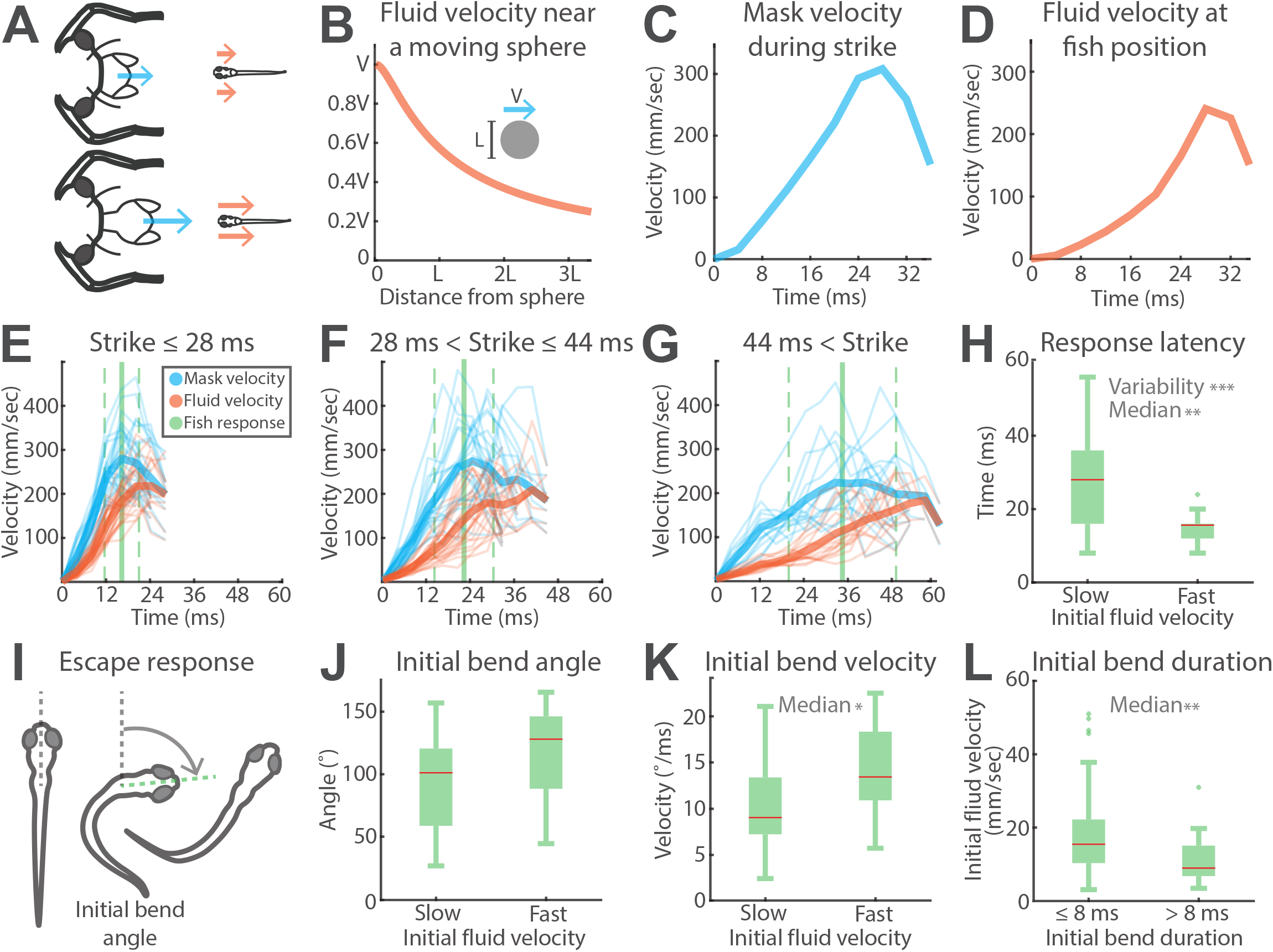
Larval zebrafish responding to fluid movement from the nymph’s strike. A) The mask velocity (in cyan) and the estimated perturbed fluid velocity at the fish position (in orange) both change over the duration the strike. B) The perturbed fluid velocity at some distance from the edge of a sphere with diameter L moving at velocity V was estimated with the analytical solution for potential flow around a sphere. The fluid velocity at the fish position depends on both the mask velocity and the distance from the mask. C) A representative example of mask velocity during an accurate strike. D) The corresponding estimated fluid velocity at the initial fish position computed from the measured mask velocity and the potential flow approximation. E–G) Mask (cyan) and fluid velocity (orange) profiles of different strikes grouped by the mask extension time with the mean *±* std of larval zebrafish response times (green solid and dashed vertical lines). Lighter lines represent individual velocity profiles while bold lines represent the mean for each group. Strike ≤ 28 ms n = 27, 28 ms < Strike ≤ 44 ms n = 24, 44 ms < Strike n = 15. H) Larval zebrafish response latencies grouped by slow (first quartile) and fast (fourth quartile) fluid velocities at the fish position 0–4 ms after onset of the attack (n = 18 for both groups). Slow initial fluid velocities produced escape responses with significantly longer latencies and more variable latencies. I) During the escape response, the fish changed heading direction with an initial bend and then swam away with propulsive swimming. The initial bend angle is the angle between the heading vector of the fish before the start of the escape response and the heading vector at the end of the initial bend. Responses where it was unclear whether the fish completed the initial bend before being captured were excluded from analysis of initial bend parameters. J) Initial bend angles of responses grouped by slow (first quartile) and fast (fourth quartile) fluid velocities at the fish position 0–4 ms after onset of the attack (n = 15 for both groups). There is no significant difference between the two groups. K) The initial bend velocity of responses grouped by slow (first quartile) and fast (fourth quartile) fluid velocities at the fish position 0–4 ms after onset of the attack (n = 15 for both groups). Initial bend velocities of responses to fast fluid velocities are significantly higher than those to slow fluid velocities. L) Fluid velocity at the fish position 0–4 ms after onset of the attack grouped by initial bend durations ≤ 8 ms and > 8 ms (≤ 8 ms n = 33, > 8 ms n = 24). Initial bend durations ≤ 8 ms occur in response to significantly higher initial fluid velocities at the fish.

**Figure 5:**
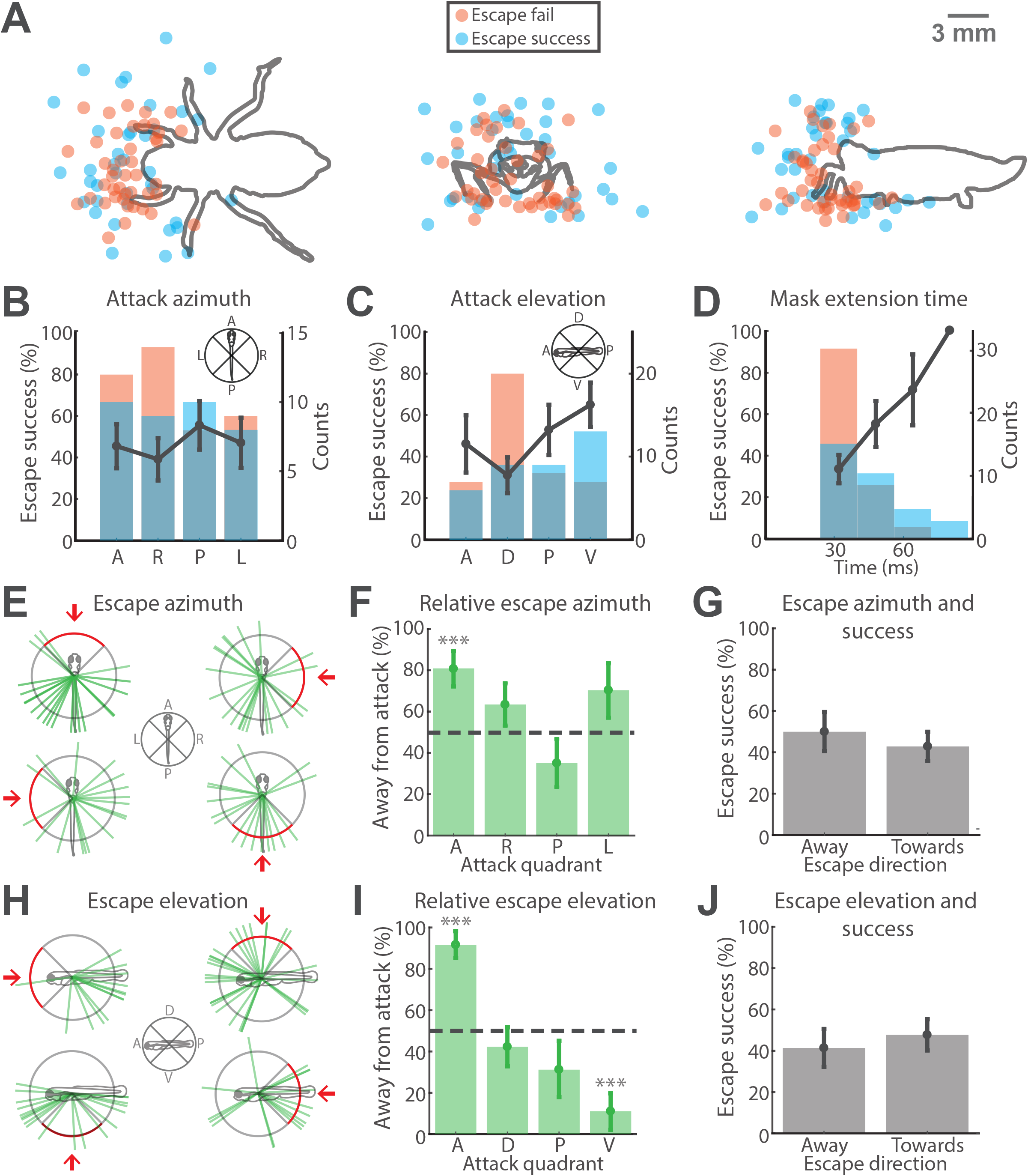
Larval zebrafish position, orientation, and escape direction. A) Top (x-y), front (y-z), and side (x-z) view orthographic projections of 3D initial fish positions before the start of an accurate strike with points colored to represent whether the fish escape response failed or succeeded in evading capture. B–D) Heights of overlapping, partially transparent bars represent the count of strikes within that bin (right y-axis) and the color represents whether the fish escape failed or succeeded. The line plot shows the mean *±* sem fish escape success probability for each bin (left y-axis, number of nymphs = 5 for all points with error bars). B) Fish escape success probability based on the azimuthal quadrant of the strike with respect to the fish. There are no significant differences in escape success probability with respect to attack azimuth. C) Fish escape success probability based on the elevation quadrant of the strike with respect to the fish. Fish escapes are significantly more likely to succeed when responding to attacks from quadrant V (ventral) than from quadrant D (dorsal). D) Histogram demonstrating the fish escape success probability based on the mask extension time to the fish position. Fish escapes are more likely to succeed as mask extension times increase. E) The azimuthal direction of fish escape represented by green lines and grouped by the azimuthal quadrant of the strike. F) Escape azimuth of fish relative to the azimuthal quadrant of the attack. The height of the bars represent the mean *±* sem probability of fish escaping away to the opposite azimuthal hemisphere from the attack quadrant (number of nymphs = 5 for all groups). Asterisks represent significant difference from 50% (dashed line). G) The azimuthal direction of fish escapes grouped by whether the response was directed towards the opposite azimuthal hemisphere or towards the same azimuthal hemisphere containing the azimuthal quadrant of attack (mean *±* sem, number of nymphs = 5 for both groups). There is no significant difference in escape success between responses with azimuthal directions towards or away from the strike azimuth. H) The elevation direction of fish escape represented by green lines and grouped by the elevation quadrant of the strike. I) Escape elevation of fish relative to the elevation quadrant of the attack (number of nymphs = 5 for all groups). Asterisks represent significant difference from 50% (dashed line). J) The elevation of fish escapes grouped by whether the response was directed away from or towards the elevation direction of the strike (mean *±* sem, number of nymphs = 5 for both groups). There is no significant difference in escape success between responses with elevation directions towards or away from the strike.

Ten different classifiers were trained for each parameter by pseudo-randomly selecting 85% of the dataset each time for training the classifier and the remaining 15% of the dataset was used for testing the accuracy of the trained classifier. This procedure allowed for an estimate of the mean *±* sem of accuracy for a classifier built on a single parameter.

To understand the interaction of parameters, 10 classifiers were also trained using all of the parameters as predictors with the same process of splitting the data as described above. The parameter importance is computed by the random forest classifier algorithm [49]. Intuitively, parameter importance is estimated through out-of-bag estimates of variable importance by permuting the values of one of the variables to see how dramatically it changes the model predictions. This random permutation is performed multiple times for each feature. The rationale behind this process of random permutation is that the most important feature will have the most negative impact on the predictions when that feature is randomly permuted across the samples. Therefore, the features are ranked by how their random permutations impacts the predictions.

Random forest classifiers were chosen over other classifiers due to their simplicity, their ability to perform nonlinear classification and exploit interactions between predictors while reducing variance through bagging/bootstrapping models. Moreover, random forest classifiers are able to provide importance values for each predictor which was especially useful for our case.

### 2.9 Computational motor volume of larval zebrafish

We computationally simulated the larval zebrafish motor volume to investigate how the intersection of the motor volume of larval zebrafish with the swept volume of the dragonfly nymph mask influenced the likelihood of survival for the fish. We used the non-engulfed fraction of the larval zebrafish motor volume as a measure of the survival likelihood of the fish. Here we describe the parameters used for the simulations.

During an escape response, larval zebrafish reorient with an initial bend with little or no movement of their center of mass and swim away with undulatory swimming during a propulsive stage [50, 51]. In this study, the fish motor volume was computationally generated to mimic this movement using a bend velocity (in 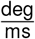) and a propulsive velocity (in 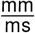). The bend velocity was first used to compute the time needed for reorientation and then the propulsive velocity was used to compute the distance traveled. In this manner, the rotation from the bend velocity and translation due to the propulsive velocity together could define the positions in 3D space that the center of mass of the larval zebrafish could reach given a certain amount of time. For all simulations of fish motor volumes, the propulsive velocity used was 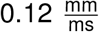 which is supported by existing literature [52, 53]. We assumed that the fish could reorient in any direction by changing pitch and yaw as necessary, which was supported by our own data (Figure 5E and H) and existing literature [50].

We did not use a full kinematics model of the fish, which would also have to account for bend acceleration and propulsive acceleration. Accounting for these accelerations would largely influence the estimates of rotation and translation at the initiation of the escape response but leave other estimates unchanged. Moreover, these accelerations are not well reported for larval zebrafish and would require imaging at far higher frame rates (> 1000 fps) to measure accurate values.

For Figures 7A and B, the fish motor volume was generated using the average bend velocity of 14 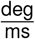 as found in this study (Figure 4K). For Figure 7C, fifty fish motor volumes were generated by pseudo-randomly sampling with replacement the initial bend velocities measured in this study (Figure 4K) for 6 increments of time remaining at escape (7, 15, 20, 25, 35, and 50 ms). These volumes were then used to compute the proportion of the fish motor volume not engulfed by the swept volume of the prehensile mask.

**Figure 6:**
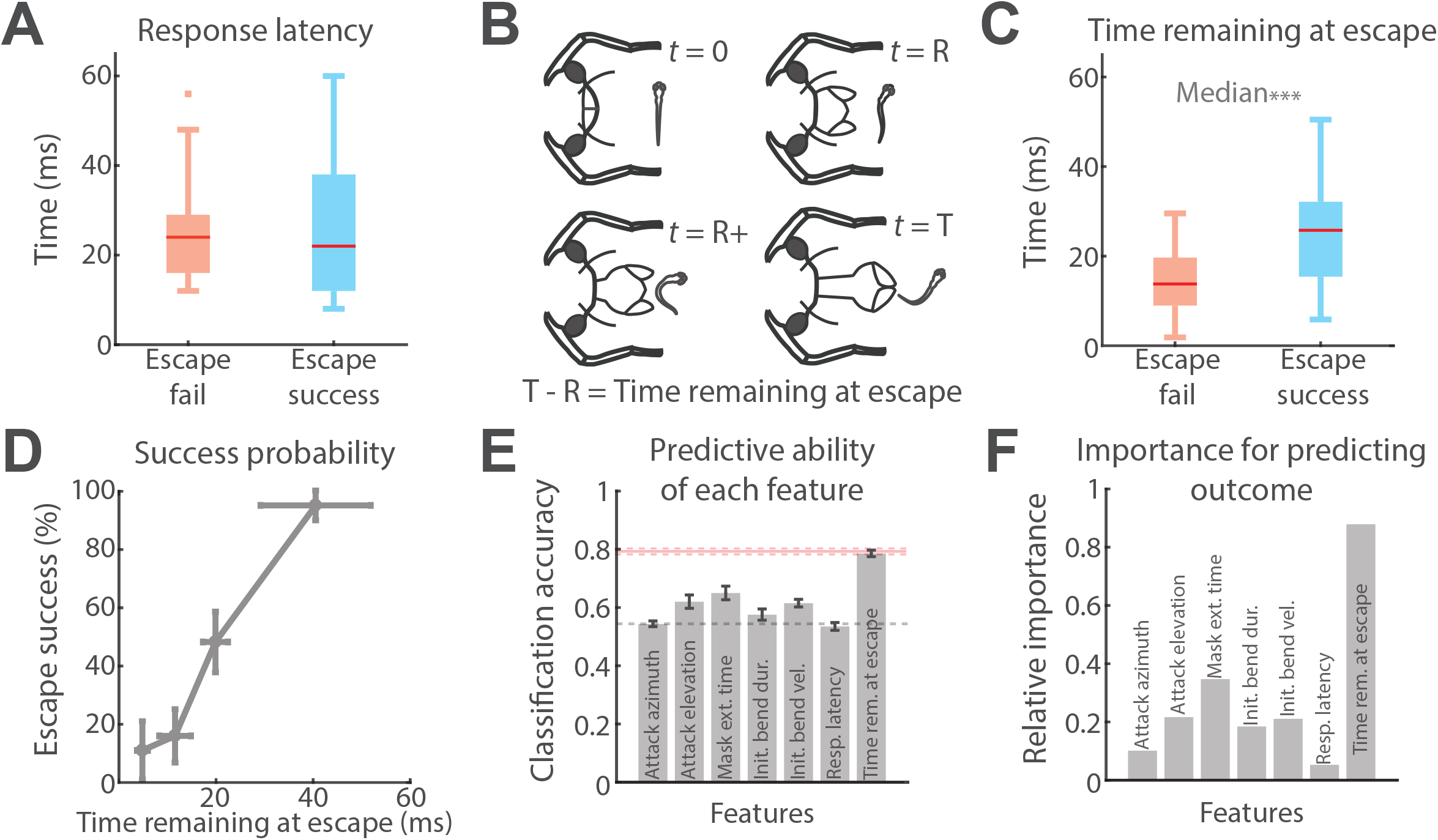
Importance of time remaining at escape. A) There is no significant difference between response latencies of failed and successful fish escapes (Escape fail n = 43, Escape success n = 37). B) The time remaining at escape is the time at the start of the fish escape response (t = R) subtracted from the mask extension time (t = T). C) The time remaining at escape is significantly longer for successful escapes than for failed escapes (Escape fail n = 43, Escape success n = 37). D) Escape success probability as a function of time remaining at escape binned into quartiles (mean *±* std). Escape success probability increases with time remaining at escape. E) Classification accuracy (mean *±* sem) of random forest classifiers in predicting escape outcome when trained on only one parameter with 10-fold cross validation. The gray dashed line is the naive classification accuracy of 0.55 and the pink dashed line at 0.79 is classification accuracy of a random forest classifier trained on all of the parameters to predict the escape outcome. The classifier using only time remaining at escape significantly outperformed classifiers trained on any of the other parameters and is not significantly different from a classifier trained on all of the parameters together. F) The relative importance of parameters in a random forest classifier trained to predict escape outcomes using all parameters. Time remaining at escape is dramatically more important than other parameters.

**Figure 7:**
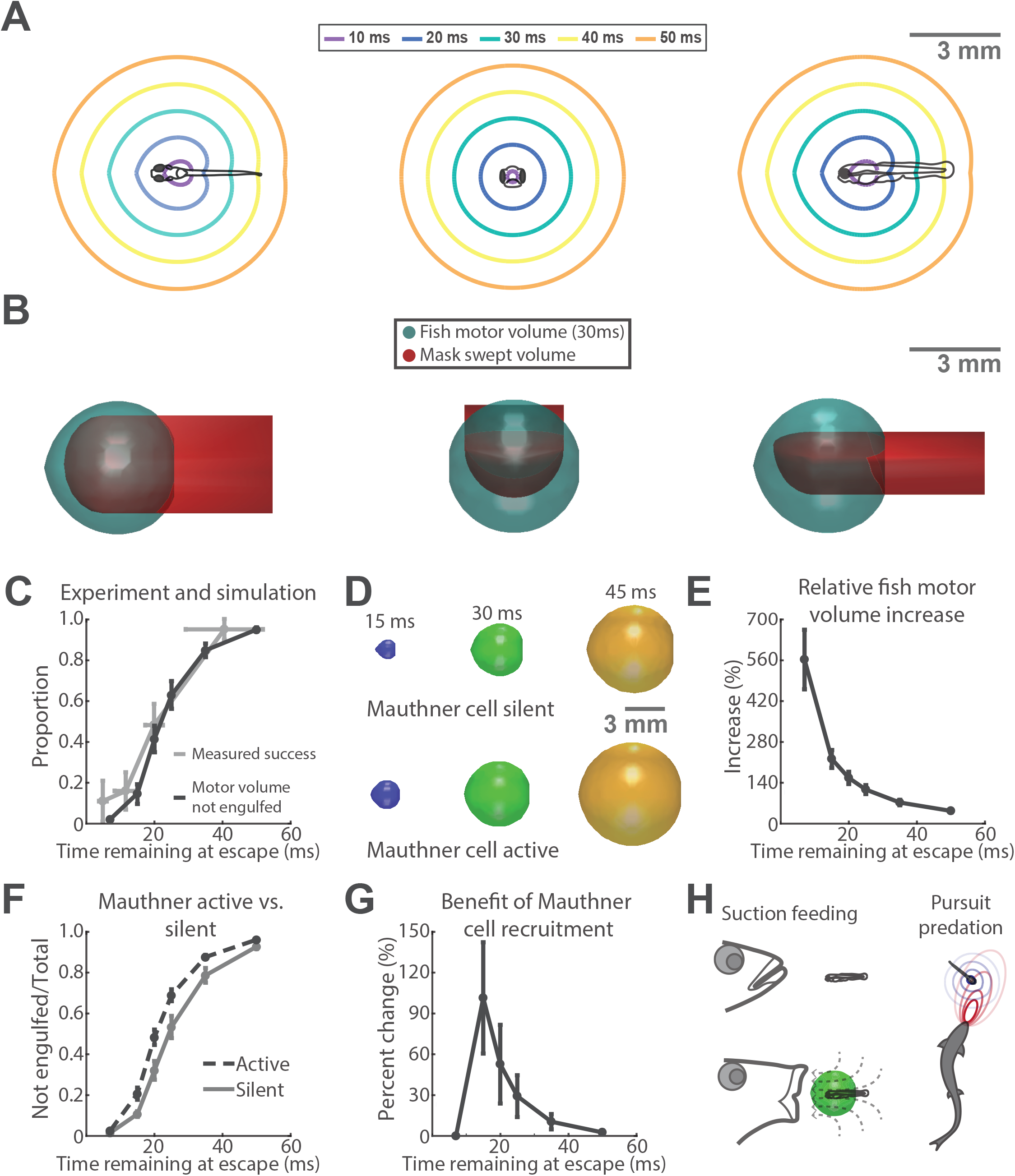
Fish motor volume in the time remaining at escape. A) Top, front, and side view cross sections of surface isochrones of the larval zebrafish motor volume for times within the range of values for time remaining at escape. This larval zebrafish volume was computationally generated using the average initial bend velocity (14°/ms) in this study and a propulsive velocity of 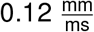. B) Top, front, and side view perspectives visualizing the intersection of the mask swept volume with the larval zebrafish motor volume for the time remaining at escape of 30 ms. C) The darker line represents the proportion of the larval zebrafish motor volume not engulfed by the mask swept volume for different values of time remaining at escape (mean *±* std). The lighter line represents the proportion of successful escape responses as a function of time remaining at escape binned into quartiles (mean *±* std). The non-engulfed fraction of the motor volume increases with time remaining at escape. There is no significant difference between the proportion of the motor volume not engulfed and the proportion of successful escape responses. D) Visualization of the estimated larval zebrafish motor volume at different times remaining at escape for the Mauthner-silent and Mauthner-active responses. The initial bend velocity used to generate the motor volumes were different for Mauthner silent 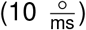 and active 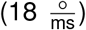 responses but the same propulsive velocity 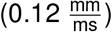 was used for both. E) The relative increase in motor volume for different values of time remaining at escape when comparing Mauthner-silent to Mauthner-active volumes (mean *±* sem). Larval zebrafish motor volumes were generated by pseudo-randomly sampling different initial bend velocities with uniform likelihood for Mauthner silent 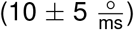 and active 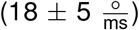 responses with 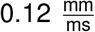 propulsive velocity for both. The increase in volume is greatest for shorter values of time remaining at escape. F) The proportion of the larval zebrafish motor volume not engulfed by the mask swept volume for different values of time remaining at escape for Mauthner silent and active responses (mean *±* std). There is a significant difference in the proportion of the fish motor volume not engulfed for Mauthner active responses. G) The percent change in the proportion of the fish motor volume not engulfed when comparing Mauthner silent to active responses. The most dramatic increases are seen for mid-range values of time remaining at escape. H) Suction feeding: larval fish motor volume and predator ingested volume interaction during suction feeding. The ingested volume is similar to mask capture volume. Pursuit predation: predator and prey motor volume interaction during pursuit predation.

The swept volume of the mask was represented by a hemi-ellipsoid (major axis: 3.2 mm and minor axis 2.8 mm) attached to the end of a half-cylinder (diameter: 3.2 mm and length: 6 mm, Figure 7B) whose dimensions were determined by measuring the prehensile masks of specimens in this study. To intersect the mask swept volume with the fish motor volume, the center of the hemi-ellipsoid of the mask swept volume was placed at the starting position of the fish motor volume. We simulated attacks from different directions with combinations of attack azimuths (front, side, and behind) and elevations (in-plane, above, and below) by rotating the mask swept volume around the origin of the fish motor volume. Ten thousand points were pseudo-randomly generated within the fish motor volume and the proportion of points not within the mask swept volume provided a measure of the proportion of the fish motor volume not engulfed.

This was carried out for each fish motor volume for each increment of time remaining at escape to find the mean *±* std proportion not engulfed for Figure 7C. Fish motor volumes representing Mauthner active and silent responses (Figure 7D) were generated by pseudo-randomly sampling different uniform distributions of initial bend velocities. The range of bend velocities for Mauthner silent motor volumes was 10 *±* 5 deg/sec and for Mauthner active motor volumes it was 18 *±* 5 deg/sec based on prior studies that investigated bend velocities of free swimming larval zebrafish before and after Mauthner ablation [41, 42, 16]. The ranges were also constructed to have some overlap since studies have found that Mauthner active and silent responses can on occasion produce similar kinematics [43, 44]. Even though there is some evidence that Mauthner cell recruitment leads to faster propulsive velocities, the simulated propulsive velocity was not changed for Mauthner active motor volumes since this effect is not clear [54, 55, 56, 53].

Intersections of the Mauthner active and silent motor volumes with the prehensile mask swept volume were carried out in the way described above. The MATLAB code used to generate and visualize larval zebrafish motor volumes and the mask swept volume can be found in the GitHub repository associated with this study.

### 2.10 Statistical methods

The statistical tests used to test hypotheses and their associated p-values are reported within the Results with references to each figure panel. The p-values are also repeated within the Results section when discussing those hypotheses and the results of the statistical test. We considered p-values < 0.05 to be significant. Within the figure panels asterisks are used to represent p-values in the corresponding manner: *p* < 0.05: *, *p* < 0.01: **, and *p* < 0.001: ***.

## 3 Results

### 3.1 Dragonfly nymph prehensile mask motor volume and attack outcome

To evaluate predator-prey interactions, we filmed strikes of nymphs upon larval zebrafish using high-speed videography at 250 frames per second (fps) with top and side view perspectives (Figure 1A). First, to quantify the biomechanical performance of the attack, we studied the duration of the predatory strike with respect to the furthest point reached by the prehensile mask in 3-dimensional (3D) space. The mask extension times ranged from 24– 176 ms depending upon the location of the strike. The time needed for the mask to reach specific positions in space (Figure 1B) was well described by a k-nearest neighbor (k-NN) model (R^2^ = 0.7, Figure 1C, see also Methods) which served as a representation of the motor volume of the mask—the volume swept by the appendage over all strikes within a given amount of time [40]. As seen in the cross-sections of the isochronic surfaces of the motor volume (Figure 1C), strikes directed towards lateral and caudal positions took more time than strikes directed medially and rostrally. This model represents the maneuverability of the mask, providing insight about the time-scale and directional bias of predatory strikes.

Next, to see how the outcome of the attack was influenced by aspects of the predatory encounter, we categorized the interactions. Successful and failed captures had distinct spatial distributions of the position of the fish before the attack (Figure 2A), where nymphs were more likely to capture larvae in closer proximity. Upon further examination, we found different kinds of failed and successful captures (Figure 2B). A failed capture could occur either due to an error in the predatory attack or an effective escape executed by the fish. A successful capture occurred either due to a fish not responding or an ineffective escape attempt by the fish. These scenarios are illustrated in Figure 2B.

The initial event is the strike by the nymph. This can lead to a failed capture when the strike is inaccurate. This happened in some instances when the fish performed spontaneous swimming movements just before the start of a predatory strike (more commonly, the fish is stationary in the period immediately preceding the initiation of the strike). In these cases, the dragonfly nymph would strike at positions where the fish was no longer present (Figure 2B, Tactile delay, Visual delay). A tactile delay refers to an inaccurate strike aimed at or near a part of the dragonfly nymph body that was touched by a spontaneously swimming fish some time before the strike began. A visual delay refers to an inaccurate strike aimed at a position previously occupied by a spontaneously swimming fish before the attack started. This suggests that the nymph is not ballistically intercepting the prey by predicting its future location, but rather striking at the position of the prey before the attack began (see Supplement). Sometimes, the nymph also made targeting errors where the fish was stationary through the entire strike but the strike was aimed inaccurately (Figure 2B, Targeting error, see Supplement). While attack errors provide information about the sensorimotor limitations of the dragonfly nymph, they cannot help determine the relevant escape decisions of the fish that confer success. Thus, we focused the remainder of our analysis on the instances where the mask was correctly aimed at the position of the fish prior to the initiation of the attack.

### 3.2 Likelihood of a response from zebrafish larvae to accurate strikes

To investigate the sensorimotor performance of the fish in this predatory context, we studied the likelihood of a fish initiating an escape response given an accurate strike. The spatial distributions of the fish positions before the start of the predatory strike were different for scenarios where the fish responded or did not respond (Figure 3A). Fish with initial positions closer to the nymph mouth were less likely to produce a response (Figure 3A). The azimuthal or elevation position of the nymph mouth with respect to the fish had little or no influence over the response probability of the fish (Figure 3B, One-way anova *p* = 0.94 and Figure C, One-way anova *p* = 0.75). However, the fish was more likely to respond given longer mask extension times (Figure 3D One-way anova, *p* = 0.01). The reduced likelihood of responding to short extension times may be because fish were captured before the initiation of a response could begin. Regardless, since the response probability is consistently above 50%, fish were always more likely to respond than not respond.

To understand whether the larval fish were initiating escape responses only on the basis of flow stimuli or a combination of visual and flow stimuli, we tested the role of flow sensing by compromising the lateral line in a group of larvae with exposure to neomycin sulphate (see Methods). This technique induces cell death in lateral line hair cells while leaving inner ear hair cells intact [46, 47]. To ensure recovery of larval zebrafish, we waited for 3 hours after neomycin treatment. Additionally, we monitored larvae to confirm that they performed spontaneous swimming movements and responded with escape maneuvers to touch stimuli before placing them in the arena with the nymph. All neomycin treated fish (n = 61) failed to respond to any strikes and were eventually successfully captured by accurate strikes (Figure 3E Mann-Whitney U test, *p* = 0.003). These data suggest that fish generate escapes in this scenario largely on the basis of flow sensing and do so with high likelihood regardless of relative orientation and position with respect to the nymph.

To better understand how the perturbed fluid movement due to the strike influenced the larval zebrafish response (Figure 4A), we next tracked the 3D position of the mask for all strikes that produced an escape response in fish. The perturbed fluid velocity in water around a moving body, such as a sphere, is a function of the velocity of the body and the distance from the body (Figure 4B, see Methods). To account for both velocity and distance when estimating the fluid flow experienced by fish, we used the mask velocity (Figure 4C) to estimate the perturbed fluid velocity over time at the initial fish position (Figure 4D). Since the distance to the initial fish position from the mask reduced over time as it extended, the perturbed fluid velocity at the initial fish position approached the velocity of the mask (Figure 4D). Different mask extension times had different velocity and associated fluid velocity profiles (Figure 4E–G). The mean response latency of the fish and the variance of response latency both increased with increasing mask extension times (Figure 4E–G). We further investigated how larval zebrafish escape responses differed in response to the earliest fluid perturbations caused by the prehensile mask. Fish responded with shorter and less variable response latencies (Mann-Whitney U test *p* = 0.002, Levene’s test, *p* = 0.0003) to the fastest quartile of fluid velocities computed between 0–4 ms after the onset of the attack (Figure 4H), hereafter referred to as the initial fluid velocity.

During the escape response, the fish changed heading direction with an initial bend and then swam away with propulsive swimming. To test whether the initial fluid velocities experienced by the fish influenced the kinematics of the escape response, we measured the bend angle (Figure 4I) for all responses where the fish clearly finished the initial bend. The bend angle was not significantly different between the fastest and slowest quartile of initial fluid velocities (Figure 4J, Mann-Whitney U test and Levene’s test, all *p* > 0.3). However, the fastest initial fluid velocities did produce responses with significantly faster bend velocities (Figure 4K, Mann-Whitney U test, *p* = 0.02). Since we recorded video at 250 fps (4 ms resolution), all initial bend durations recorded fell between 4– 8 ms (n = 33), 8–12 ms (n = 22), or 12–16 ms (n = 2) after the onset of the escape. Escapes with shorter bend durations (≤ 8 ms) tended to be in response to significantly higher initial fluid velocities (Figure 4L, Mann-Whitney U test, *p* = 0.008). These data suggest that fish modulate their escape responses by deploying maneuvers with different latencies and kinematics based on the magnitude of the perturbed fluid velocity.

### 3.3 Larval zebrafish position, orientation, and escape direction

We next examined how the position and orientation of the fish at the start of the attack, along with its escape direction, influenced escape success. The spatial distribution of the fish positions before the start of the predatory strike for successful and failed escapes largely overlapped but failed escapes tended to happen closer to the nymph (Figure 5A). The azimuthal quadrant of the attack with respect to the fish had no significant influence on escape success probability (Figure 5B). However, the attack elevation was significantly related to the escape success probability (One-way anova, *p* = 0.001). Specifically, fish were significantly more likely to execute successful escapes when responding to attacks from below (ventral, V) than from above (dorsal, D, Figure 5C, Mann-Whitney U test, *p* = 0.005). Moreover, fish were more likely to execute successful escape maneuvers in response to longer mask extension times (Figure 5D, One-way anova *p* < 0.001), which is consistent with the higher probability of escaping at further distances or with more time.

We then analyzed escape direction with respect to the attack to determine whether fish escaped away from the attack and how escape direction influenced escape success. Escape directions in the opposite hemisphere of the attack quadrant were considered to be away from the attack. When escape directions were grouped by the azimuthal or elevation attack quadrants (Figure 5E, H), fish did not consistently move away from the attacks (One-sample proportion tests). Escapes in response to attacks from the right (R), posterior (P), and left (L) azimuthal quadrants were not significantly biased away from the attack (Figure 5F). Similarly, escape directions in response to attacks from the dorsal (D) and posterior (P) elevation quadrants were not significantly biased away from the attack (Figure 5I). While fish did significantly bias their escape directions away from attacks in the anterior (A) azimuthal and elevation quadrants, this could be because escape movements typically involve a turn [44], especially those in response to attacks directed at the head [57]. The lack of consistent directional control was also illustrated by the fact that larvae would often escape in the direction of attacks occurring in the ventral (V) elevation quadrant (Figure 5I). Critically, whether fish escaped away or towards the attack had no significant influence on escape success probability (Figure 5G, J, Mann-Whitney U Tests, all *p* > 0.5). However, escape trajectories toward the attack can occur along pitch or yaw angles that take the fish around the mask, thereby keeping the fish out of the capture zone.

Together these data suggest that the attack azimuth and the escape direction relative to the attack direction were not significantly related to escape success probability and, in some cases, escapes toward the attack were successful. Thus it seems more important that zebrafish move quickly rather than in the opposite direction.

### 3.4 Importance of time remaining at escape

Having found that the escape direction of the larval fish had no discernible influence on escape success, we next explored how the fish’s response latency may have influenced the escape outcome. Surprisingly, the response latencies of successful and failed attacks were not significantly different (Figure 6A, Mann-Whitney U test, Levene’s test, all *p* > 0.8). To investigate how this discrepancy might be explained, we examined in more detail how response latency may interact with mask extension time. To do so, we defined the time left from the initiation of the fish escape response until the mask reaches the initial position of the fish as the time remaining at escape (Figure 6B). The time remaining at escape was dramatically different for successful and failed escapes (Figure 6C, Mann-Whitney U test, *p* ≪ 0.0001). Additionally, the escape success probability increased with increasing time remaining at escape (Figure 6D, One-way anova *p* ≪ 0.001).

To further investigate the interaction of escape maneuver parameters and which single parameter, if any, had the most influence on the outcome we trained different random forest classifiers with each parameter to predict escape success or failure (see Methods). The parameters were the following: 1) attack azimuth; 2) attack elevation; 3) mask extension time; 4) bend duration; 5) bend velocity; 6) response latency; 7) time remaining at escape. Parameters known not to be related to escape outcome (attack azimuth and response latency) were included as controls along with kinematic parameters (bend duration and bend velocity) which had not yet been tested. Ten different classifiers were trained for each parameter by selecting 85% of the dataset (n = 68) each time for training and testing accuracy with the remaining 15% of the dataset (n = 12). This allowed for an estimate of the mean *±* sem of accuracy for a classifier trained on each parameter.

A naive estimator which predicted that all escapes failed had a classification accuracy of 0.55 (grey dashed line Figure 6E). Any classifier trained on one of these parameters with a significantly higher classification accuracy suggested that the parameter had some influence on the outcome. The classifiers trained on attack elevation, mask extension time, bend velocity, and time remaining at escape had classification accuracies significantly different from the naive estimate. However, the classifier trained on time remaining at escape dramatically outperformed all other classifiers (pairwise Mann-Whitney U tests with Bonferroni correction, all *p* < 0.005) and was not significantly different from a classifier trained on all of the parameters together (pink dashed line Figure 6E, Mann-Whitney U test, *p* = 0.4). As a further test, a random forest classifier trained on all of the parameters together was used to determine which parameter was the most important in determining the model prediction (see Methods). The importance of time remaining at escape as a parameter was dramatically higher than all other parameters in determining the model prediction (Figure 6E).

These data suggest that even though various parameters were correlated with escape outcome, the time remaining at escape was the best and most important predictor of the escape outcome (Figure 6E and F).

### 3.5 Fish motor volume in the time remaining at escape

The time remaining at escape limits the volume of space that contains all possible trajectories of the fish. This constraint is visible in the cross-sections of isochronic surfaces representing the fish motor volume for different times remaining at escape (Figure 7A). These isochrones quantify the maneuverability of the fish [40] within that time. The visualization of the fish motor volume in Figure 7A was generated computationally (see Methods) using the average bend velocity measured in this study (14°/ms) during the initial bend and the average swimming velocity during propulsion (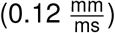) measured in this study of and confirmed by others [52, 53]. Using this model, we visualized the portions of the fish motor volume not engulfed by the swept volume of the mask (Figure 7B). The non-engulfed fraction of the fish motor volume represents the regions of space visited during a successful escape. We hypothesized that the importance of the time remaining at escape in determining the outcome was due to its direct influence on the fish motor volume and its intersection with the mask swept volume.

We used simulations to investigate how the fraction of the fish motor volume not engulfed by the mask volume corresponded to the escape success probability (Figure 6D). Fifty virtual larval zebrafish motor volumes were generated by pseudo-randomly sampling initial bend velocities measured in this study and using a propulsive velocity of 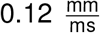 for values of time remaining at escape from 7–50 ms. These virtual motor volumes were intersected with virtual mask swept volumes attacking from different directions with respect to the fish to measure the proportion of the motor volume not engulfed by the mask (see Methods). The increase in the proportion of the fish motor volume not engulfed with increasing time remaining at escape corresponded well to the increase in the measured escape success probability (Figure 7C). Moreover, an analysis of covariance demonstrated that the fraction of the motor volume not engulfed and the proportion of successful escape responses as a function of the time remaining at escape were not significantly different (slope *p* = 0.4, intercept *p* = 0.2).

We used the proportion of the fish motor volume not engulfed to computationally investigate the utility of recruiting the Mauthner cell in generating a response. Since the recruitment of the Mauthner cell is known to produce the fastest bend velocities [41, 42, 44], we generated virtual Mauthner-active and Mauthner-silent fish motor volumes using different ranges of initial bend velocities but the same propulsive velocity, as used earlier (Methods). The virtual Mauthner-active motor volumes were consistently larger than the virtual Mauthner-silent motor volumes for all times remaining at escape (Figure 7D). The greatest difference between the Mauthner-active and Mauthner-silent motor volumes were seen for the shortest times remaining at escape (Figure 7E). This is because the parameters for the Mauthner-active motor volume allowed the virtual fish to finish the initial bend earlier and start propelling away, which more rapidly increases the size of the motor volume.

Mauthner-active and Mauthner-silent motor volumes were also intersected with the swept volume of the nymph mask to compute the proportion of the fish motor volume not engulfed (Figure 7F, refer Methods). The proportion of the Mauthner-active fish motor volume not engulfed was significantly different from the proportion of the Mauthner-silent motor volume not engulfed (Two-way anova, Mauthner recruitment p < 0.001, Time remaining at escape p < 0.001, Interaction p = 0.2).

To quantify the benefit of Mauthner cell recruitment at different times remaining at escape, we calculated the percent change in the proportion of the motor volume not engulfed from Mauthner-silent to Mauthner-active volumes (Figure 7G). Our simulations showed that Mauthner activation increased the proportion of the fish motor volume not engulfed by the mask by 30–100% on average for times remaining at escape of 15–25 ms. Times remaining at escape longer than 40 ms showed a reduced benefit of Mauthner cell recruitment since slower escapes would be equally effective in evading the mask’s swept volume. Moreover, times remaining at escape shorter than 7 ms also had reduced benefit from recruiting the Mauthner cell since that amount of time is inadequate for the fish to move out of the mask swept volume.

This analysis demonstrates how simulations of prey motor volume and predator swept volume can be used to estimate the utility of recruiting specialized escape circuits in response to an attack. Moreover, the intersection of these volumes provides insight into how the time remaining at escape shapes the outcome of the predatory interaction. Also, it may clarify the lack of impact of response latency on escape success, since similar response latency values can be associated with different values of time remaining at escape.

## 4 Discussion

Our goal was to evaluate how escape maneuver parameters of prey interact in order to influence survival by analyzing the escape responses of larval zebrafish to attacks from dragonfly nymphs. We identify the time remaining for the dragonfly mask to reach the fish from the onset of the fish’s escape response as the most predictive parameter for escape success. We call this parameter the time remaining at escape and explain its role in determining the volume of space that contains all possible trajectories of the fish—the fish motor volume. Using a computational approach, we estimate the fish motor volume for different times remaining at escape to quantify the fish’s ability to evade the capture volume of the nymph—the volume swept by the mask. Additionally, we use this approach of analyzing motor volumes to calculate the utility of recruiting the Mauthner neuron for generating the escape by estimating the relative increase in escape success probability.

We argue that the time remaining at escape robustly determines the outcome since it serves as a limiting constraint on the possible trajectories of the fish, as visualized by the time-limited fish motor volume [40]. This perspective can clarify the influence of response latency, speed, and direction of an escape maneuver on evasion success across different predation contexts. For the same time remaining at escape, faster escape speeds would increase the size of the fish motor volume and therefore, increase the proportion of the motor volume not engulfed by the capture volume of the predator. This explains the existing evidence in support of the benefit of fast speeds during escape [20, 22]. However, escape responses to slower predators that leave more time remaining at escape may not require fast escape speeds for successful evasion. This explains the evidence found in other studies against the need for fast escape speeds [23, 24, 50].

Similarly, shorter escape response latencies for the same dragonfly mask extension times would increase the time remaining at escape. This would also increase the fish motor volume and thus, the non-engulfed fraction. Unexpectedly, we find that the response latency of larval fish was not significantly different for failed and successful escapes. We argue that this is due to the variability of dragonfly mask extension times (Figure 5D) where the same fish response latency can produce a successful escape in the case of longer extension time and a failed escape in the case of the shorter extension time. However, for scenarios where the duration of predatory strikes are more consistent, changes in the latency of escapes would produce measurable changes in evasion success, as seen in other studies [23, 5, 25, 58].

The response latency of the prey animal is also related to the distance at which the prey can sense the predator. Sensing range has been found to be key to evasion success in some studies [25, 59]. While this parameter may be more relevant to pursuit predation in fish than to ambush predation, it can still be understood within the framework of motor volumes and the time remaining at escape. Clearly, with longer sensing range, the prey is afforded more time remaining at escape, increasing its motor volume and chances of surviving a predatory strike. However, it’s important to note that escaping earlier can also allow the predator more opportunity to change course and follow suit. In these scenarios, the prey may wait to initiate an evasive maneuver even after seeing the predator so that it can outmaneuver the predator at closer distances [60] where the motor volume of the slower or stopped prey is less directionally biased than that of the fast moving predator.

Finally, specific escape directions that lead the prey out of the capture volume of a predator would also lead to successful escapes. In our study, for longer times remaining at escape, we find that a number of directions in the nearly spherical motor volume of the fish led out of the capture volume of the dragonfly nymph. This result aligns with studies which find that successful escape trajectories are not required to follow a single optimal trajectory [11] and can even be directed towards the attack [61]. However, scenarios where the predator capture volume engulfs a large portion of the prey motor volume may leave only a subset of directions that successfully evade the attack. For such cases, the appropriate choice of escape direction would be vital to survival, as shown in modeling studies [3, 6] and behavioral experiments [62]. In our data, when the mask engulfed a large portion of the fish motor volume for shorter times remaining at escape, fish had low survival rates possibly because their escapes were not directed away from the attacks.

Different predatory scenarios may change the relative importance of response latency, speed, direction, or even other parameters in producing successful escapes. However, here we unify the influence of these parameters on the escape outcome by clarifying their role in the generation of the critical non-overlapping regions of the predator and prey volumes.

We found that escapes were initiated in response to the flow stimulus of the mask extension as suggested by the dramatic reduction of response after neomycin treatment of fish. Even though the fluid flow caused by mask movement was critical in the initiation of the escape response, fish did not consistently escape away from the direction of the attack. The lack of correlation between the attack direction and escape direction aligns with existing findings of larval zebrafish escapes initiated by flow sensing [63].

However, our findings do indicate that the magnitude of the fluid perturbation influenced the escape response. Faster fluid velocities due to mask movement produced shorter latency escape responses, with faster initial bends, and shorter initial bend durations. These escape kinematics suggest that fish were more likely to recruit the Mauthner cell in response to higher magnitude fluid perturbations. The argument for differential Mauthner cell recruitment based on stimulus parameters is well supported by previous findings which show that fish perform a graded assessment of threat [44].

Using simulations of fish motor volume and mask swept volume intersections, we were also able to estimate the utility of recruiting the Mauthner cell for an escape maneuver. Since the recruitment of the Mauthner cell generates responses with the highest initial bend velocities [41, 42, 44], we compared Mauthner-active with Mauthner-silent motor volumes generated with different bend velocities. Our simulations showed that Mauthner activation dramatically increased the proportion of the fish motor volume not engulfed by the mask for a specific range of times remaining at escape from 15–25 ms. Since this range composes a significant proportion of the experimentally observed range of times remaining at escape, there is a clear functional benefit of recruiting the Mauthner neuron in this predatory context. This is further supported by recent experimental evidence which shows that larval zebrafish are far less likely to survive dragonfly nymph strikes after Mauthner cell ablation [16].

However, our simulations also suggested a reduced benefit from recruiting the Mauthner cell for larger values of time remaining at escape because motor networks producing slower movements would be equally effective. This result aligns with existing findings which demonstrate that fish are less likely to deploy a Mauthner active escape in response to slower approaching predators that will take longer to reach the fish [64, 44]. Surprisingly, there was also reduced benefit from recruiting the Mauthner cell for very small values of time remaining at escape since these values do not allow for enough time to maneuver out of the predator capture volume. This potentially explains freezing behavior in cases where there is not enough time to escape [65, 66, 67, 44].

Our results extend directly to aquatic suction feeding (Figure 7H) since the volume ingested by the predator is analogous to the mask swept volume. Moreover, the ingested volume changes in time [68] creating a flow stimulus that initiates the escape response [69] and leaves some time remaining after escape until the total volume is ingested. In support, other researchers have demonstrated that the accuracy of aiming this ingested volume during suction feeding is critical in determining outcome of the strike [70]. Poorly aimed ingested volumes during suction feeding would overlap less with the prey motor volume leaving regions of safety for the prey to escape into. The prevalence of suction feeding in a variety of fish [71] indicates that the framework presented here fits a wide array of aquatic predatory interactions.

For prey, the time remaining at escape is related to the speed of the predatory attack at the moment the attack is sensed. Though it is unclear whether animals estimate this parameter for flow stimuli, studies using looming objects suggest that animals do estimate the time remaining to capture for visual stimuli [72, 73, 74]. Conceivably, faster attacks that produce more intense sensory stimuli push the estimates of time remaining to lower values. These estimates of time remaining directly correspond to the utility of deploying different escape maneuvers. This aligns with existing evidence of more intense stimuli producing shorter latency and higher speed escape responses in other animals [12, 75]. Given the importance of time remaining in predicting the escape outcome, the evidence that prey estimate this parameter, and its ability to determine the utility of different escape responses, we expect that time remaining at escape is a major driver of decision-making and a source of significant selection pressure. The larval zebrafish is an ideal model system to pursue how the computation of the time remaining at escape may be implemented by the nervous system.

More generally, comparing predator and prey motor volumes provides a method to quantify the maneuverability of each agent through the predatory interaction. This also applies to pursuit predation (Figure 7H) where the motor volumes change over time as predator and prey attempt to outrun and out-maneuver each other [76, 60]. The regions of the prey motor volume not intersecting with the predator engulfing volume denote the subset of movements and corresponding neural circuits that constitute successful evasive strategies. This subset has clear implications for decision making during escape and the evolutionary pressure on the selection of appropriate maneuvers to increase survival. The approach presented here provides a method to estimate the utility of specific escape maneuvers by connecting the interplay of many temporal and kinematic parameters to their influence in shaping the reachable spaces of predator and prey.

## A Supplement

### A.1 Dragonfly nymph predatory performance

On occasion, the dragonfly nymph made targeting errors when attacking the larval zebrafish (Figure A1A) and would strike at a position where the fish was never present. These targeting errors were more likely when the fish was within a certain region of the visual field of the nymph (Figure A1B, C) which has implications for the visual acuity or motor-coordination of the dragonfly nymph. This region of the visual field corresponding to a higher likelihood of targeting errors was below the center-line of the dragonfly nymph’s eyes (Figure A1D). Interestingly, the dragonfly nymphs were also more likely to make targeting errors when the fish faced away from the nymph (Figure A1E, F). This may be due to the lack of the stark visual contrast afforded by the dark, black eyes of the larval zebrafish. Finally, longer mask extension times when the fish was at further distances from the nymph were also more likely to cause targeting errors (Figure A1G).

Beyond targeting errors, the dragonfly nymph also made other errors when hunting the larval zebrafish. These errors were sometimes due to visual and tactile delays. In these cases, the dragonfly nymph would strike at positions where the fish was no longer present. In the case of a tactile delay, the fish would swim by and graze or nearly-graze the body of the nymph and the nymph would strike at the position near its body even though the fish was no longer there when the strike began. In the case of the visual delay, the fish would swim through the visual field of the dragonfly nymph (not near its body) and the nymph would strike at a point along the fish’s trajectory of motion but the fish was not there when the strike started. Each case allowed for a measurement of the sensorimotor delays of the nymph by measuring the time from when the fish last inhabited the targeted position to the initiation of the strike. Sensorimotor delays of the nymph for tactile stimuli: 75.5 *±* 30 ms (n = 16); for visual stimuli: 278 *±* 110 ms (n = 5).

## B Acknowledgments

We thank Neelesh Pantankar for his guidance with fluid mechanics. Additionally, we acknowledge members of the McLean lab for their advice in experimental protocols and Elissa Szuter for technical help maintaining the fish colony. This work was supported by NSF-IOS 1456830 and NIH R01-NS067299.

---

**SM Figure A1:**
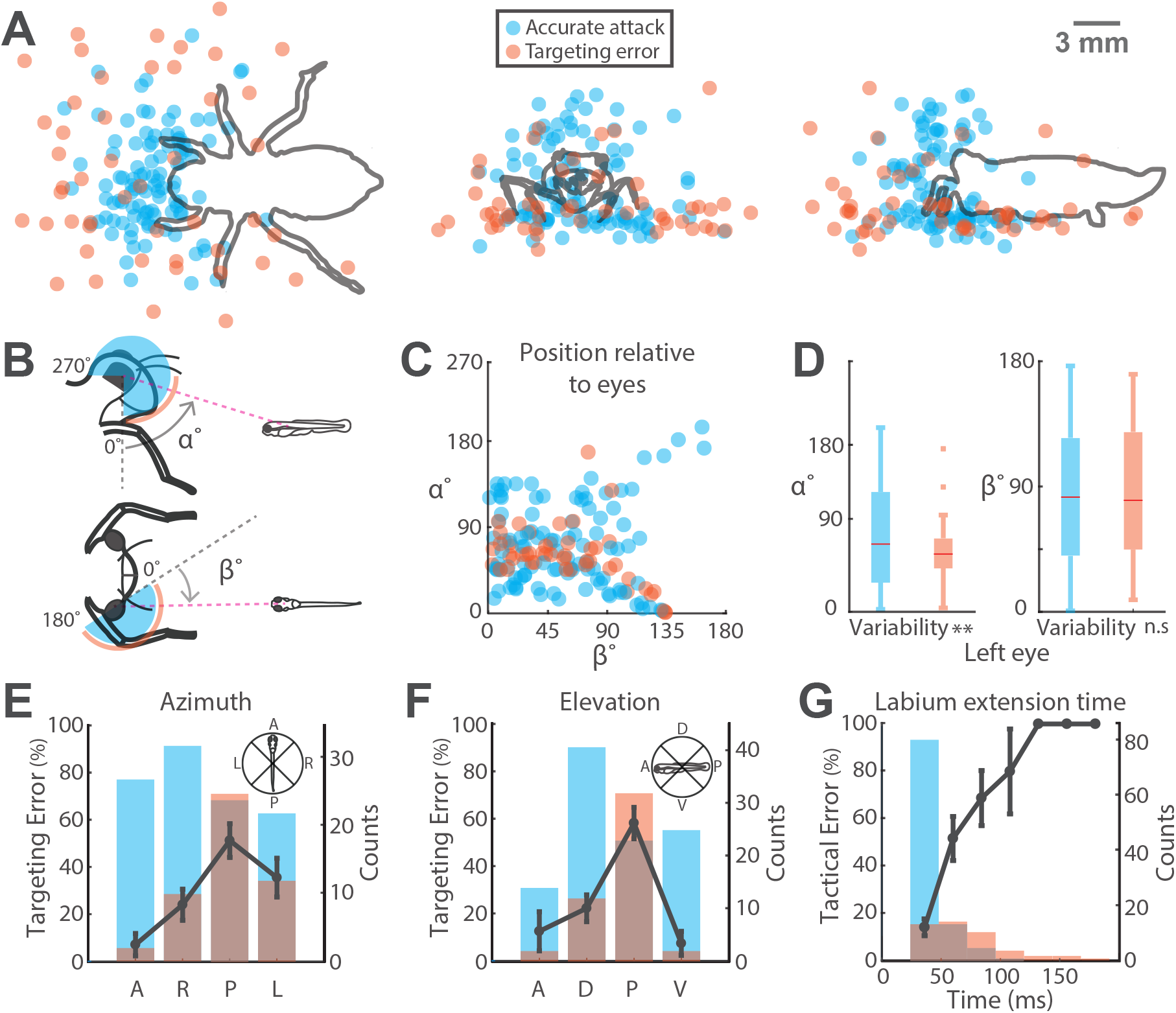
A) Top (x-y), front (y-z), and side (x-z) view orthographic projections of 3D initial fish positions before the start of the labial strike with points colored to represent an accurate attack or a targeting error by the nymph. Legend provided in the figure. B) Diagram describing angles *α* and *β* measuring the fish swim bladder position with respect to the nymph eye. C) Scatter plots of *α* and *β* on both eyes for fish positions not in physical contact with the nymph (number of nymphs = 5, accurate attack n = 65, targeting error n = 28). D) Box-and-whisker plots demonstrating the distribution of *α* and *β* values for accurate attacks and targeting errors just for both eyes. Accurate attacks and targeting errors have significantly different variability of *α* values (Levene’s test p = 0.01) but no significant difference in the medians. There is no significant difference in *β* values for accurate attacks and targeting errors. E) Targeting error probability based on the azimuthal position of the nymph head with respect to the fish. Azimuthal positions of the nymph were grouped by quadrants shown in the panel. Heights of each bar represent the count of labial strikes within that quadrant (right y-axis) and the color represents accurate attacks or targeting errors. The line plot shows the mean *±* sem targeting error probability for each quadrant (left y-axis, number of nymphs = 5 for all quadrants). Nymphs are significantly more likely to make a targeting error when in azimuthal quadrant P and significantly less likely to make a targeting error when in azimuthal quadrant A (one-way anova: p < 0.001, pairwise Mann-Whitney U tests with Bonferroni correction: all p < 0.01). F) Targeting error probability based on the elevation of the nymph head with respect to the fish. Elevation positions of the nymph were grouped by quadrants shown in the panel. Heights of each bar represent the count of labial strikes within that quadrant (right y-axis) and the color represents accurate attacks or targeting errors. The line plot shows the mean *±* sem targeting error probability for each quadrant (left y-axis, number of nymphs = 5 for all quadrants). Nymphs are significantly more likely to make a targeting error when in elevation quadrant P (one-way anova: p < 0.001, pairwise Mann-Whitney U tests with Bonferroni correct: all p < 0.001). G) Histogram demonstrating the probability of the nymph making a targeting error based on the labium extension time to the fish position. Measured labium extension times are shown for accurate attacks but labium extension times for targeting errors were estimated with the labial motor volume model. Heights of each bar represent the count of labial strikes within that bin (right y-axis) and the color represents accurate attacks or targeting errors. The line plot shows the mean *±* sem targeting error probability for each bin (left y-axis, number of nymphs = 5 for all points with error bars). Nymphs are significantly more likely to make a targeting error for labial strikes longer than 44ms (one-way anova: p < 0.001, pairwise Mann-Whitney U tests with Bonferroni correction: all p < 0.01).

## Notes

### Competing Interest Statement

The authors have declared no competing interest.

